# Postmortem Cortex Samples Identify Distinct Molecular Subtypes of ALS: Retrotransposon Activation, Oxidative Stress, and Activated Glia

**DOI:** 10.1101/574509

**Authors:** Oliver H. Tam, Nikolay V. Rozhkov, Regina Shaw, Duyang Kim, Isabel Hubbard, Samantha Fennessey, Nadia Propp, The NYGC ALS Consortium, Delphine Fagegaltier, Lyle W. Ostrow, Hemali Phatnani, John Ravits, Josh Dubnau, Molly Gale Hammell

**Author notes:** These authors contributed equally to this work (Oliver H. Tam, Nikolay V. Rozhkov, Regina Shaw).

## Abstract

Amyotrophic Lateral Sclerosis (ALS) is a fatal neurodegenerative disease characterized by the progressive loss of motor neurons. While several inherited pathogenic mutations have been identified as causative, the vast majority of cases are sporadic with no family history of disease. Thus, for the majority of ALS cases, a specific causal abnormality is not known and the disease may be a product of multiple inter-related pathways contributing to varying degrees in different ALS patients. Using unsupervised machine learning algorithms, we stratified the transcriptomes of 148 ALS decedent cortex tissue samples into three distinct and robust molecular subtypes. The largest cluster, identified in 61% of patient samples, displayed hallmarks of oxidative and proteotoxic stress. Another 20% of the ALS patient samples exhibited high levels of retrotransposon expression and other signatures of TDP-43 dysfunction. Finally, a third group showed predominant signatures of glial activation (19%). Together these results demonstrate that at least three distinct molecular signatures contribute to ALS disease. While multiple dysregulated components and pathways comprising these clusters have previously been implicated in ALS pathogenesis, unbiased analysis of this large survey demonstrated that sporadic ALS patient tissues can be segregated into distinct molecular subsets.

## Introduction

ALS is a fatal progressive neurodegenerative disorder with no known cure, and only two FDA-approved treatments that appear to mildly slow disease progression. ALS is a largely sporadic disease, with 90% of patients carrying no known genetic mutation or family history of disease. Large-scale patient sequencing studies have identified a growing number of genes in which mutations are linked to ALS (Chia et al., 2018; Nicolas et al., 2018; van Rheenen et al., 2016). The most common ALS-associated mutations are repeat expansions in the intronic region of C9orf72, while mutations in well-known ALS-associated genes such as SOD1 and TDP-43 are typically present in fewer than 2% of all ALS patients (Chia et al., 2018).

Mutations in the TDP-43 gene are rare in ALS, yet nearly all ALS patients exhibit cytoplasmic aggregates of TDP-43 in the affected tissues (Arai et al., 2006; Neumann et al., 2006). TDP-43 has known roles in RNA splicing, stability, and small RNA biogenesis (Cohen et al., 2011). Recently, several studies have suggested that TDP-43 also plays a role in regulating the activity of retrotransposons (Krug et al., 2017; Wenxue Li et al., 2015; Saldi et al., 2014). Retrotransposons, a subset of transposable elements (TEs), are genomic parasites capable of inserting new copies of themselves throughout the genome by a process called retrotransposition. Previous work from our lab and others has shown that TDP-43 represses retrotransposon transcripts at the RNA level in animal models of TDP-43 pathology (Krug et al., 2017; Wanhe Li et al., 2012). However, a role for TDP-43 in general retrotransposon silencing has not been demonstrated, nor whether TDP-43 pathology in ALS patients correlates with retrotransposon de-silencing. Of note, prior studies have identified a link between retrotransposon expression and repeat expansion in another ALS-linked gene, C9orf72 (Prudencio et al., 2017). Finally, contrasting studies either failed to find an enrichment for elevated levels of the endogenous retrovirus HERV-K in a smaller sample of ALS tissues (Mayer et al., 2018) or suggested that TDP-43 may activate HERV-K transcription rather than silencing this particular retrotransposon (Wenxue Li et al., 2015). These studies left open the question of whether retrotransposon silencing is a conserved role for TDP-43 and whether retrotransposon de-silencing would be expected in human tissues with TDP-43 dysfunction.

Here we show that robust retrotransposon de-silencing occurs in a distinct subset of ALS patient samples, and that this is associated with TDP-43 dysfunction. Novel unbiased machine learning algorithms identified three distinct ALS patient molecular subtypes within the large ongoing sequencing survey by the NYGC ALS Consortium. The largest subgroup of patients (61%) showed evidence of oxidative and proteotoxic stress. A second subgroup (19%) displayed strong signatures of glial activation and inflammation. A third subgroup (20%) was marked by retrotransposon re-activation as a dominant feature. Given the novelty of retrotransposon expression as a feature of some ALS patient samples, we explored the mechanistic connection to TDP-43 associated pathways and TDP-43 pathology for these samples in a smaller validation cohort. These subtypes may reflect different predominant aberrant cellular mechanisms contributing to motor neuron cell death in ALS pathogenesis, and thus suggest that specific therapeutic strategies may have relevance to distinct subgroups of sporadic ALS patients.

## Results

### Evidence for Distinct Molecular Subtypes in ALS Patient Samples

The NYGC ALS Consortium has gathered deeply sequenced transcriptomes from the frontal cortex of 77 ALS patients as well as 18 Neurological and Non-Neurological controls (Fig 1A). For some patients, multiple samples were taken from various regions of the frontal cortex, including motor cortex, such that 148 total transcriptomes were available from ALS patients, while 28 were from controls (176 samples in all) (See also Supp. Table S1). Most of these patients presented with sporadic ALS disease, i.e. no known family history or pathogenic mutation, consistent with general estimates that ALS as a disease is largely sporadic (Chia et al., 2018).

**Fig. 1.**
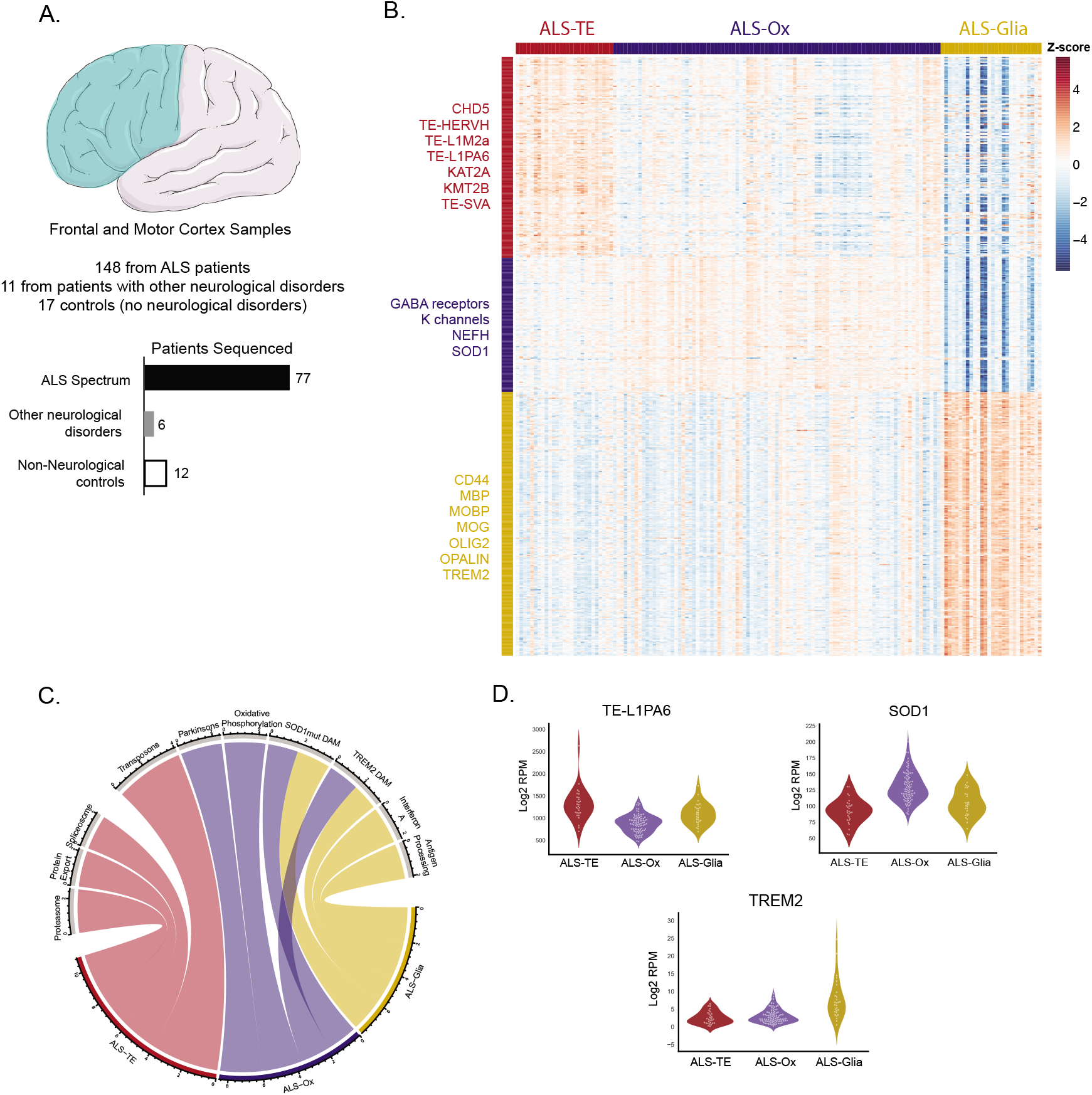
ALS Patients in the NYGC ALS Consortium Cohort show 3 distinct subtypes: Retrotransposon Reactivated (ALS-TE), Oxidative Stress Signatures (ALS-Ox), and Glial Dysfunction (ALS-Glial). **(A)** A schematic depiction of the frontal and motor cortex regions from which post-mortem tissue samples were obtained: 148 from ALS patients, 11 from patients with other neurological disorders, and 17 from control subjects with no sign of neurodegenerative disease. Multiple samples were obtained for most patients, such that these samples represent 77 ALS patients, 6 patients with other neurological disorders, and 12 healthy controls. **(B)** De novo clustering of transcriptomes from the 148 ALS patient samples using NMF-based algorithms returned 3 robust and distinct groups. The retrotransposon reactivated set showing high TE expression as a dominant signature formed its own subgroup (ALS-TE), representing 20% of all patient samples. Two additional groups were identified which showed signatures of oxidative stress (ALS-Ox, 61% of all samples), and glial dysfunction (ALS-Glial, 19% of samples). Representative genes and transposon markers returned as marking each subset are identified along the left of the heatmap. **(C)** Gene Set Enrichment Analysis was performed separately for each group of patients (ALS-TE, ALS-Ox, and ALS-Glial) vs. Controls, respectively. Patients in the ALS-TE group showed strong enrichment for all Expressed Transposons as well as depletion for genes involved in Spliceosomal components, Proteasomal components, and Protein Export Factors. ALS-Ox patients showed strong enrichment for genes involved in Oxidative Phosphorylation as well as oxidative stress genes linked to Parkinson’s Disorder. ALS-Glia patients showed inflammatory pathways linked to Interferon α and Antigen presentation as well as pathway previously identified as upregulated in Disease Associated Microglia (DAM) induced via SOD1 or TREM2 mutations. **(D)** Violin plots of example markers for each group show the LINE element L1PA6 marking the ALS-TE group, SOD1 marking the ALS-Ox group, and TREM2 marking the ALS-Glial samples.

The large size of this dataset enables *de novo* clustering algorithms to determine whether the ALS patient samples fall into distinct molecular subsets defined by specific gene signatures. We used a recent method for *de novo* clustering that was originally developed for single-cell sequencing datasets, but has been equally validated on bulk transcriptomes, SAKE (Ho et al., 2018). SAKE is based on non-negative matrix factorization (NMF) and can robustly estimate both the number of clusters present in a given dataset and the confidence in assigning each sample to a cluster. SAKE returned optimal results for 3 clusters within the NYGC ALS Consortium Cohort (Supp. Fig S1), and a heatmap of the genes that define these clusters is given in Fig 1B. Based on the gene markers returned by the SAKE algorithms, some of which are marked on the left hand side of Fig 1B (See also Supp. Table S2), we have chosen to re-label the 3 ALS subtypes as ALS-TE (elevated Transposable Element, or TE, expression, red in Fig 1B), ALS-Ox (oxidative stress markers, blue), and ALS-Glia (elevated glial markers, gold).

The largest subset of patient samples in the NYGC survey (91/148) were marked by elevated levels of several genes previously associated with ALS. These included NEFH, SOD1 and CDH13 (Chia et al., 2018), as well as a general expression signature consistent with oxidative and proteotoxic stress. These samples are identified as ALS-Ox (Fig 1B, blue). ALS-Ox patient samples showed elevated levels of genes in the Oxidative Phosphorylation pathway, in pathways associated with Parkinson’s Disease, as well as proteotoxic stress pathways as determined by GSEA (Fig 1C, Supp. Table S3). Surprisingly, these samples also showed enrichment for genes previously noted to be elevated in both mutant SOD1 (Chiu et al., 2013) and TREM2-dependent (Keren-Shaul et al., 2017) models of disease associated microglia (Fig 1C), though microglial markers were not specifically enriched in these samples (Supp. Table S2).

A second subset of patient samples, dubbed ALS-Glial (Fig 1B, gold), is defined by increased expression of genes that mark astrocytes (CD44, GFAP) and oligodendrocytes (MOG, OLIG2) suggesting that glial markers were a dominant signature in the transcriptomes of these samples. The ALS-Glial subset of patient samples also showed strong enrichment for transcriptional signatures previously identified as marking disease-associated microglia in either a mutant SOD1 (Chiu et al., 2013) or TREM2-dependent model (Keren-Shaul et al., 2017) (Fig 1C, see also Supp. Table S4) and in levels of TREM2 expression (Fig 1D). Finally, these patients also showed upregulation of innate immune pathways that are typically elevated in activated microglia, such as Interferon and antigen processing pathways (Fig 1C).

Retrotransposon transcripts form a subset of the genes that define the remaining ALS samples, ALS-TE, named for transposable elements (TEs). These patient samples (Fig 1B, red) represented 20% (29/148) of the samples in the NYGC cohort. The ALS-TE samples showed an enrichment for transposon expression as the most significantly enriched pathway relative to controls using GSEA pathway analysis (Fig 1C, Supp. Table S5), and this included TEs from the LINE, SINE, and LTR classes. The samples in the ALS-TE subset also showed depletion for components of the spliceosome and for protein export pathways, pathways previously linked to normal TDP-43 function (Cohen et al., 2011). Finally, violin plots of individual transposable elements, such as the LINE element L1PA6, show increased transposon levels for specific retrotransposons (Fig 1D, Supp. Table S5). The fact that unbiased and unsupervised clustering identified several individual TEs as specifically defining this group provides a rigorous and quantitative method of determining the samples in which TE expression is well above the normal control levels and unique to this subset of ALS patients.

As stated above, multiple frontal and motor cortex tissues were sampled for a subset of ALS patients, allowing for an estimation of the level of concordance of ALS subtype between different tissues for the same patient. Of the 40 patients with both frontal and motor cortex samples sequenced, we note that only 7 were discordant between the two tissues (17.5%), suggesting a high degree of overlap between these two tissues (Supp Table S6).

### The ALS-Ox Group Displays evidence of Oxidative Stress

The importance of oxidative stress in neurodegenerative disease began with the initial identification of superoxide dismutase 1 (SOD1) as the first ALS-associated mutation (Rosen et al., 1993). Subsequent studies have identified a diverse array of functional pathways altered in SOD1 mutant mouse models, including SOD-1 mediated autophagy (Rudnick et al., 2017), proteotoxic stress (Bruijn et al., 1998), and neuroinflammation (Chiu et al., 2013), reflecting both the pleiotropic roles played by SOD1 protein as well as different manifestations depending on the cellular context in which SOD1 is dysfunctional. Since that initial discovery, several genes with roles in oxidative stress, proteotoxic stress, and autophagy have been linked to ALS (Taylor et al., 2016). Consistent with the importance of these pathways, and their linked nature in generating neuronal stress, we found that 61% of the NYGC ALS patient samples displayed gene expression signatures consistent with a robust response to oxidative and proteotoxic stress, as described below.

Samples in the ALS-Ox group show elevated levels of several stress response genes, mutations in which have been linked to ALS, including SOD1 itself, as well as the neurofilament protein NEFL (Fig 2A-B). More generally the pathways that are elevated in the ALS-Ox samples relative to controls include genes involved in oxidative phosphorylation, proteasomal components, and genes involved in the unfolded protein response (Fig 2C). This subset of patients also displayed elevated expression of several genes that have been previously associated with multiple neurodegenerative diseases including both ALS and Parkinson’s Disease, such that these disease pathways are also significantly enriched in the ALS-Ox group relative to controls (Fig 2C, Supp. Table S3).

**Fig. 2.**
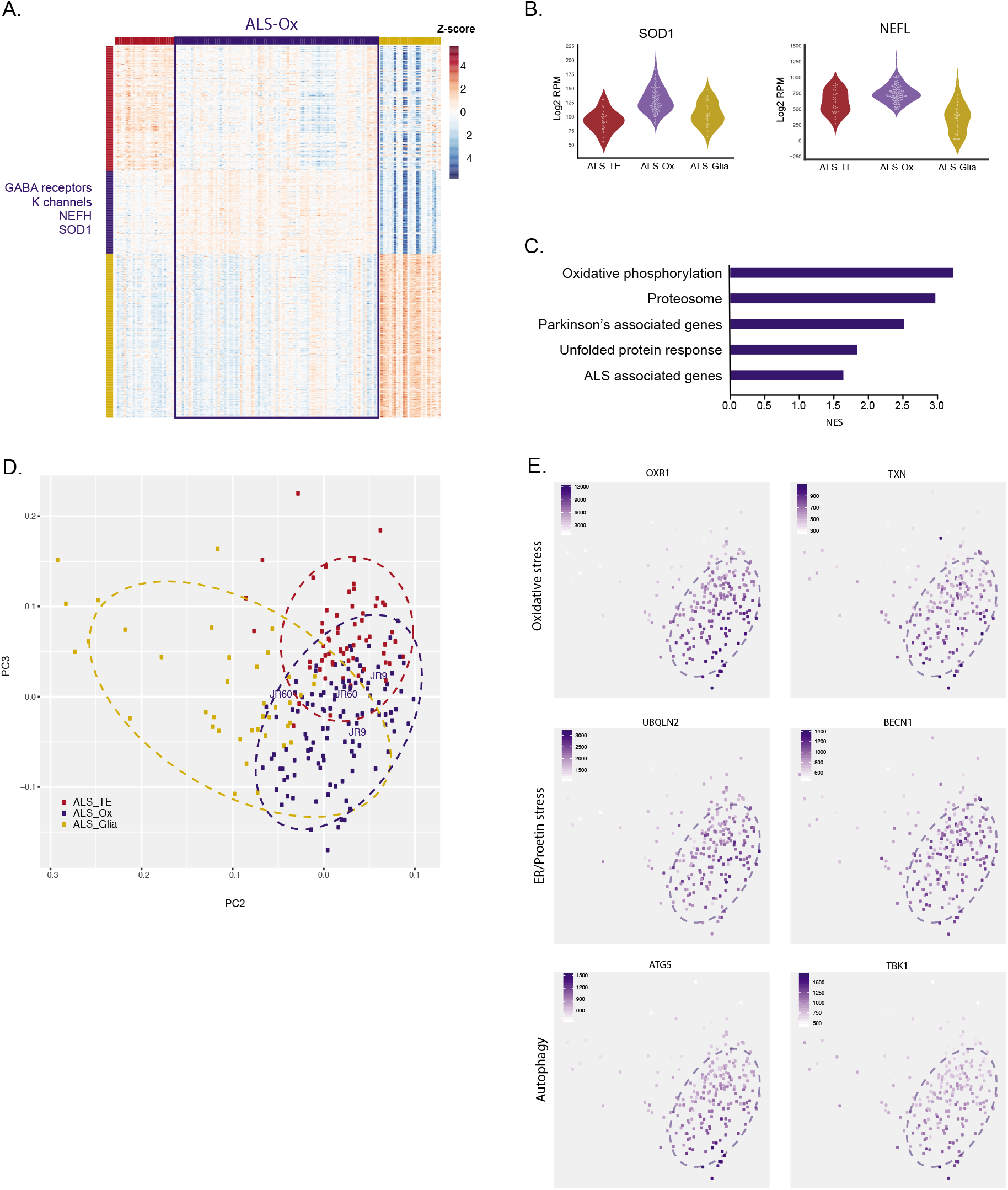
ALS-Ox Patients in the NYGC ALS Consortium show markers of oxidative stress. (A) The ALS-Ox group represents the middle 61% of patient samples in the ALS Consortium set (outlined in Blue), with ALS-Ox marker genes listed along the left hand side of the heatmap. (B) Violin plots of example markers for this group show two known ALS-associated genes, SOD1 and NEFL, specifically elevated in the ALS-Ox group (blue violins). (C) The top enriched pathways identified by GSEA in the ALS-Ox group samples include the Oxidative phosphorylation pathway, known genes associated with Parkinson’s Disease as well as ALS, and components of the unfolded protein response. (D) Additional ALS patient samples from UCSD are assigned to the NYGC ALS subgroups by comparing their similarity to each ALS Subtype, as displayed on this PCA plot (UCSD samples marked by patient ID in blue). (E) To confirm the link between ALS-Ox membership and stress pathways, individual PCA plots are shown where each sample is colored by the normalized expression of genes in the Oxidative Stress pathway (OXR1, TXN), ER-linked Proteotoxic stress pathways (UBQLN2, BECN1), and downstream genes related to autophagy (ATG5, TBK1).

To further validate these results, we obtained fresh frozen motor cortex tissue for an additional set of 13 ALS patients and 6 non-neurological controls, provided by a Tissue Bank at UCSD (Supp. Fig S2, Supp. Table S7). Two biological replicate transcriptomes were sequenced from each patient and control tissue sample to ensure the results would be robust to within-patient heterogeneity. We verified that these patient datasets displayed similar groupings to those from the NYGC cohort and quantitatively assigned these samples to each of the ALS Subtype groups, by combining the two groups into a single principal components analysis (PCA, Fig 2D). Patients were assigned to a subtype based on the distance to each cluster centroid on the PCA graph, with most samples falling within the 95% confidence interval ellipse defining the NYGC group (Supp. Fig S2). Two of the 13 ALS patients from this UCSD cohort fell within the ALS-Ox group, with 2 samples plotted per patient to show concordance between biological replicates. Consistent with their classification, we noted that these patient samples also displayed similar expression patterns for all ALS-Ox marker genes, comparable to that seen in the NYGC samples (Supp. Fig S2, grey bars). To demonstrate that position on this PCA plot represents the expression of genes underlying the oxidative and proteotoxic stress responses in both patient groups, we generated individual PCA plots for several genes marking these key stress response pathways, with the color intensity of each dot representing the expression of these marker genes in Fig 2E. For the oxidative stress pathway, expression is shown for Oxidation Resistance 1 (OXR1) and Thioredoxin (TXN). For the proteotoxic stress response pathway, expression is shown for Ubiquilin-2 (UBQLN2) and Beclin-1 (BECN1). Finally, for markers of autophagy, expression is shown for Autophagy Gene 5 (ATG5) and TANK binding kinase (TBK1). We note that these 3 pathways are typically linked, such that both oxidative stress and proteotoxic stress have been noted to induce an autophagic response in neurodegenerative disease (Wong and Holzbaur, 2015), consistent with these pathways showing elevation in the same subset of samples. Thus, these results are consistent with ALS patient samples in the ALS-Ox group mounting a robust response to several neuronal stressors, which may be concurrent in the tissue.

### The Predominance of Glial markers in the ALS-Glia Group

Several recent studies have noted the important role that glial cells play in neurodegenerative disease. Astrocytes, cells that support neuronal function by providing nutrients and removing waste, have previously been shown to secrete neurotoxic factors when expressing ALS-associated mutations (Di Giorgio et al., 2007; Nagai et al., 2007). Microglia, the innate immune cells of the central nervous system, have been shown to become activated in several neurodegenerative diseases (Deczkowska et al., 2018), setting off a neuroinflammatory cascade that eventually results in motor neuron cell death. In the ALS Consortium Cohort, 19% of the ALS patient samples showed extensive evidence of glial involvement, as discussed below.

Samples in the ALS-Glial subgroup show elevated expression of markers for all glial cell types, including Astrocytes (CD44, GFAP), Microglia (IBA1/AIF1, TREM2), and Oligodendrocytes (OLIG1, OLIG2, MOG), as noted on the heatmap of Fig 3A. In particular, some of the microglial markers most prominently elevated in these samples represent genes known to be expressed in activated microglia and associated with neurodegenerative disease, such as TREM2 and IBA1, as seen in the violin plots of Fig 3B. Consistent with this, pathway analysis of the genes that are up-regulated in ALS-Glial samples relative to controls include mediators of the inflammatory response in general, the interferon response in particular, and other genes downstream of the TNF-alpha signaling pathway (Fig 3C). Two recent reports have identified gene expression signatures of disease associated microglia (DAM) in mouse models of neurodegeneration carrying mutations in either SOD1 (Chiu et al., 2013) or TREM2 (Keren-Shaul et al., 2017). These two DAM gene expression signatures were also strongly enriched in the ALS-Glia cortex samples relative to controls (Fig 3C, Supp Table S4).

**Fig. 3.**
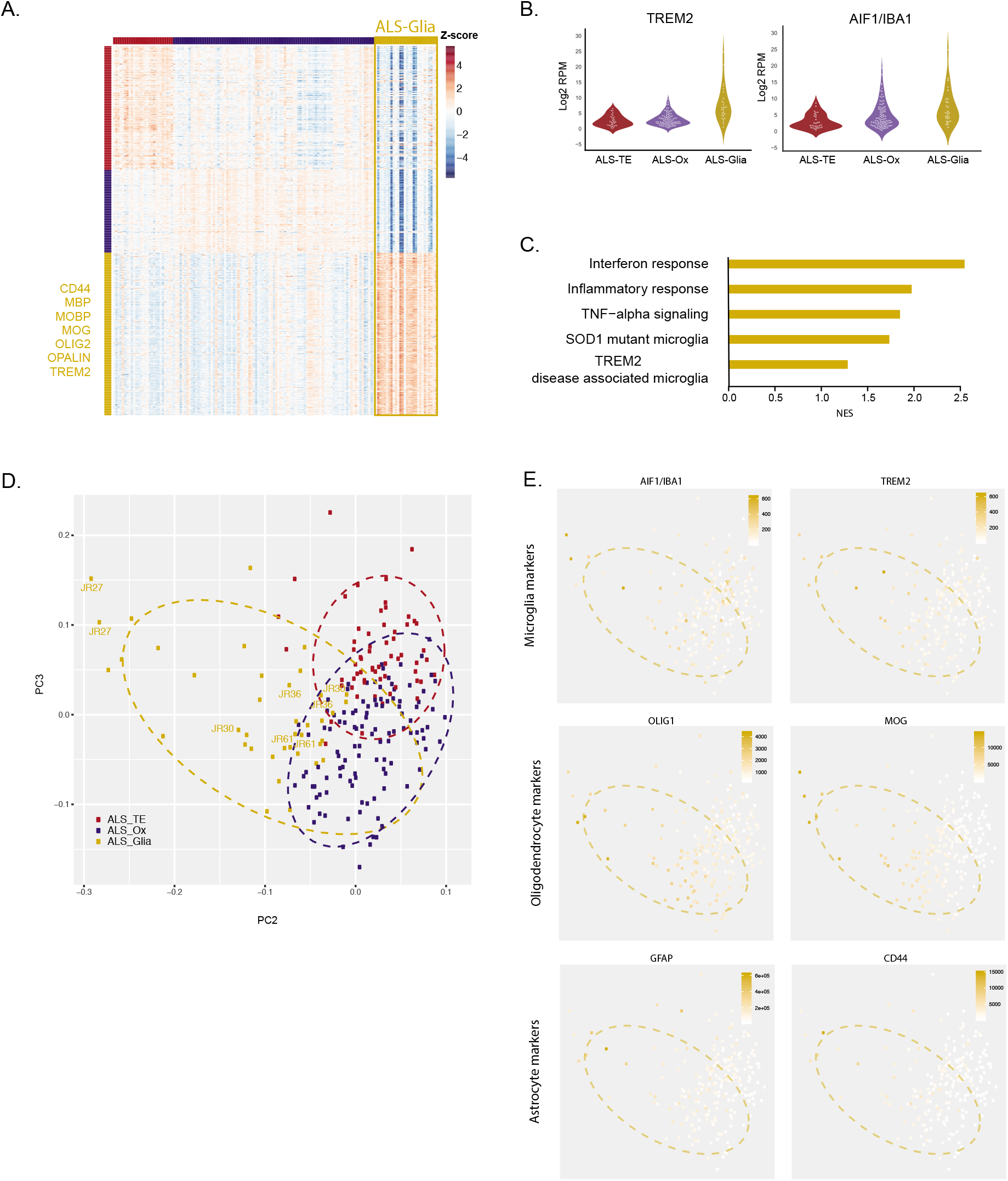
ALS-Glia Patients in the NYGC ALS Consortium Cohort show evidence of disease associated microglia. (A) The ALS-Glia group represents the 19% of patient samples on the right side of the ALS Consortium set (outlined in Gold), with additional ALS-Glia marker genes listed along the left side of the heatmap (B) Violin plots of example markers for this group show two known microglial marker genes, TREM2 and IBA1, specifically elevated in the ALS-Glia group (gold violins). (C) The top enriched pathways identified by GSEA in the ALS-Glia group samples include genes involved in the Interferon, Inflammatory response, and TNF-alpha pathways, all of which can generally be grouped as “inflammatory.” In addition, two recently identified signatures of activated microglia in SOD1 and TREM2 mutant mice were also elevated in this group. (D) Additional ALS patient samples from UCSD are assigned to the NYGC ALS subgroups by comparing their similarity to each ALS Subtype, as displayed on this PCA plot (UCSD samples marked by patient ID in gold). (E) To confirm the link between ALS-Glia membership and markers of particular glial subsets, individual PCA plots are shown where each sample is colored by the normalized expression of genes that mark Microglia (IBA1, TREM2), Oligodendrocytes (OLIG1, MOG), and astrocytes (GFAP, CD44).

One underlying mechanism that could explain profiles in the ALS-Glia samples is a relative loss of motor neurons, and/or increased presence of glial cells in these particular tissues. We used software designed to infer the relative composition of different cell types in bulk tissues, NeuroExpresso (Mancarci et al., 2017) to determine whether there was any evidence of differential cell type enrichment in the ALS-Glial samples. The NeuroExpresso results supported this model, with estimates of cellular composition showing an increase in markers of activated microglia (P<0.003) as well as oligodendrocytes (P<0.002) and astrocytes (P<0.02) (Supp. Fig 4) relative to controls. These patients also showed the strongest signatures for relative loss of neuronal markers from both the Pyramidal (P<6.7e-7) and GABAergic (P<3.1e-5) classes. We cannot differentiate whether these represent tissues with more glial activation/ proliferation/recruitment to the area, more severe motor neuron loss (and thus relative enrichment of non-neuronal signatures), or both. However, these patient tissues clearly show much stronger glial involvement relative to all other ALS patient samples and controls. No other ALS subtypes show significant differences in cell type enrichment compared to controls (Supp. Fig S3).

In our UCSD patient tissue validation cohort, 4/13 ALS patients clustered with the NYGC ALS-Glial group, without strong evidence of transposon de-silencing or oxidative stress markers (gold markers labeled by patient ID in the PCA plot of Fig 3D). These samples also showed elevated levels of TREM2 and IBA1 expression, at levels similar to that in the NYGC samples (Fig 3E). Moreover, markers of oligodendrocytes were also strongly enriched in this group, including OLIG1 and MOG. Finally, astrocyte markers were also present in these ALS-Glia samples, but were largely absent from samples in the ALS-TE and ALS-Ox groups, with GFAP and CD44 shown as representatives (Fig 3E).

### The ALS-TE Group Displays altered expression of genes in multiple pathways linked to TDP-43

Sporadic ALS patients are known to show cytoplasmic accumulation and aggregation of TDP-43 protein in the motor cortex and spinal cord, two tissues where motor neuron loss occurs (Arai et al., 2006; Neumann et al., 2006). Such TDP-43 pathology is thought to cause both a loss of the nuclear function of TDP-43 as well as aggregation associated impact. Thus TDP-43 targets would be expected to be mis-regulated in ALS patients with TDP-43 pathology and an associated loss of TDP-43 nuclear function. Previous studies have linked TDP-43 to roles in splicing, mRNA metabolism, and protein export (Cohen et al., 2011), as well as retrotransposon silencing (Krug et al., 2017; Wanhe Li et al., 2012). Consistent with this last role, several retrotransposons from the ERV, LINE, and SVA class were selected as specifically marking the ALS-TE group (Fig 4A). High levels of individual retrotransposons, including TEs from the LINE class (Fig 4B), characterize this group, while retrotransposons in general formed the top enriched pathway in GSEA (Fig 4C). Additional pathways depleted from the ALS-TE group are also consistent with previously identified functional roles for TDP-43, including protein export, the spliceosome, and the proteasome (Fig 4C). Finally, we note that TDP-43 expression levels are the lowest in the ALS-TE group overall (Fig 4B), which could be mediated by TDP-43 auto-regulation (Ayala et al., 2011) or other mechanisms.

**Fig. 4.**
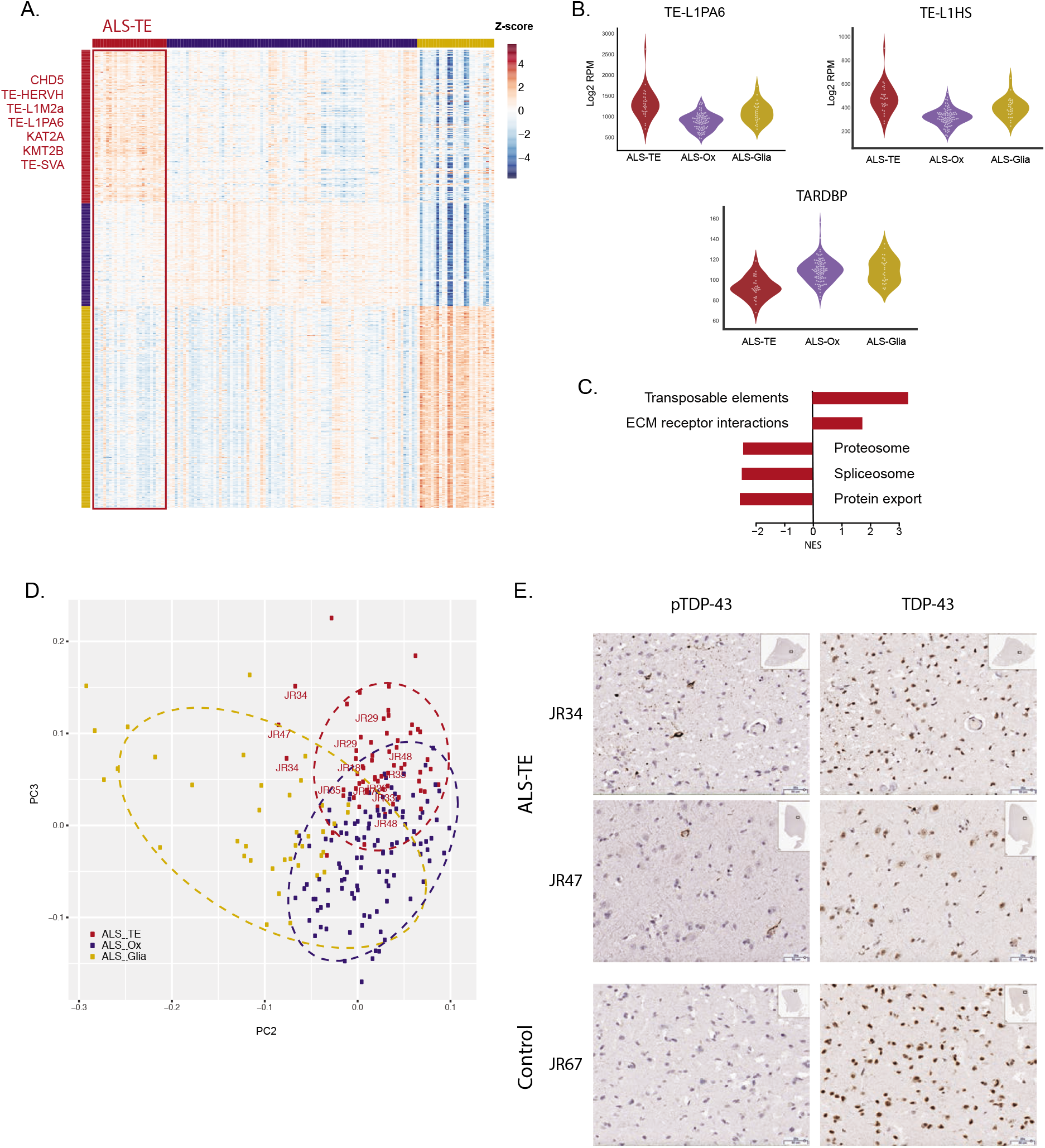
ALS-TE Patients in the NYGC ALS Consortium Cohort show retrotransposon re-activation. (A) The ALS-TE group represents the left hand 20% of patients in the ALS Consortium sample set (outlined in Red), with ALS-TE markers listed along the left hand panel (B) Violin plots of example markers for this group show two relatively young members of the LINE family of retrotransposons, L1PA6 and L1HS. Violin plots of TARDBP expression show reduced expression in ALS-TE compared to other subtypes. (C) The top enriched pathways identified by GSEA in the ALS-TE group include all expressed transposons and extracellular matrix (ECM) interactions. Genes that show the greatest depletion in these samples include members of the proteasome, spliceosome, and protein export factors. (D) Additional ALS samples from UCSD (2 replicates per patient) are assigned to the defined subgroups by comparing their similarity to each ALS Subtype, as displayed on this PCA plot (UCSD samples marked by patient ID in red). (E) To confirm the link between ALS-TE membership and TDP pathology, UCSD patient samples were stained with antibodies to the phosphorylated form of TDP-43 (pTDP-43, left) as well as with antibodies that recognize full length TDP-43 protein (right). ALS-TE samples show evidence of pTDP-43 pathology not present in Controls or other ALS subgroups.

To further validate these results, we again turned to samples from the UCSD patient cohort for follow up analysis and validation. This allowed for the identification of 7 ALS patients from the UCSD cohort that fell within the ALS-TE group, with patient ID marked in red on Fig 4D. Consistent with their PCA-based classification, we noted that these UCSD patient samples also displayed elevated levels of TEs, comparable to that seen in the NYGC samples (Supp. Fig S2). To establish a more direct link between the ALS-TE group and TDP-43, we stained representative samples from the ALS-TE group with an antibody directed against phosphorylated TDP-43, which has been previously characterized to identify patients with TDP-43 pathology in ALS (Neumann et al., 2009). The ALS-TE patients with elevated TE levels and loss of known TDP-43 pathway genes are the most likely to show TDP-43 inclusion pathology in the motor cortex, as shown for representative fields in Fig 4E. Non-ALS-TE patients and control samples did not show evidence of pTDP-43 inclusions in the tissues analyzed (Supp. Fig 4), nor did they show transcriptional patterns consistent with established TDP-43 regulated pathways (Supp Tables S3-5). Together these results establish a correlation between TDP-43 dysfunction and transcriptional re-activation of retrotransposon sequences in ALS patient tissues. The mechanistic basis for this will be explored next.

### TDP-43 Functions to Silence Retrotransposons

While pathways linked to oxidative stress and activated glia have previously been linked to ALS and other neurodegenerative diseases (Taylor et al., 2016), a potential role for retrotransposons in ALS is only recently emerging. As such, we used a genomics approach in neuronal-like cell lines to bolster the connection between retrotransposon expression and the function of the ALS associated protein TDP-43. TDP-43 is an RNA binding protein with two RNA recognition domains that recognize UGUGU repeat motifs present in thousands of cellular RNAs (Polymenidou et al., 2011; Tollervey et al., 2011). Previous TDP-43 studies using fly (Krug et al., 2017), (Wanhe Li et al., 2012), and human (Wanhe Li et al., 2012) models to identify downstream targets of TDP-43 have suggested that retrotransposons may form a subset of TDP-43 targets, either directly or indirectly, but the extent and functional impact of this was not fully understood. To establish the subset of direct TDP-43 target genes and retrotransposons, we sequenced the RNAs bound to TDP-43 protein using an enhanced cross-linking and immunoprecipitation protocol (eCLIP-seq) (Van Nostrand et al., 2016) in human SH-SY5Y neuroblastoma cells. Peaks were called using a CLIP-seq analysis tool designed for handling repetitive reads, CLAM (Z. Zhang and Xing, 2017). Results from two biological replicates of TDP-43 eCLIP libraries were merged and normalized to a crosslinked, size-matched input control, resulting in 36,716 called peaks mapping to 5770 genes and 439 transposable elements (Fig. 5, Supp. Table S8-9). Thirty one percent of all peaks mapped to transposable elements generally (Fig. 5B). However, 58% of the TE associated peaks mapped to the opposite strand, suggesting the TEs were providing regulatory sequence for the host gene, a phenomenon previously observed for other RNA binding proteins (Attig et al., 2018; Kelley et al., 2014; Zarnack et al., 2013). The remaining 42%, which were directly bound to TEs in the sense orientation, can be grouped into three major families, encompassing LINE elements (5% of all called peaks), SINE elements (4% of peaks), and LTR regions that include endogenous retroviruses (2%). Verification that these sequences were directly bound to TDP-43 protein was supported by the presence of TDP-43 binding motifs at called peaks in both the gene and retrotransposon annotated peaks (Fig. 5C, see also Supp. Table S10). Examples of TDP-43 CLIP reads over a known gene target (TARDBP/TDP-43) as well as novel retrotransposon targets from each major class (LINE – L1PA6, SINE – AluY, LTR – HERV3, and SVA – SVA_D) can be seen in Fig 5A and 5D with a full list in Supp. Table S9. The gene targets correlate well with previously identified TDP-43 targets from CLIP-seq and RIP-seq studies (Colombrita et al., 2012; Polymenidou et al., 2011; Tollervey et al., 2011; Van Nostrand et al., 2016; Xiao et al., 2011), which previously missed transposon targets due to masking of repetitive regions or discarding of ambiguously mapped reads. Together, these results demonstrate that TDP-43 normally binds broadly to both gene and TE-derived transcripts. This comprehensive list of direct targets in SH-SY5Y cells provides a platform to investigate the impact of TDP-43 knock down on expression of target RNAs.

**Fig. 5.**
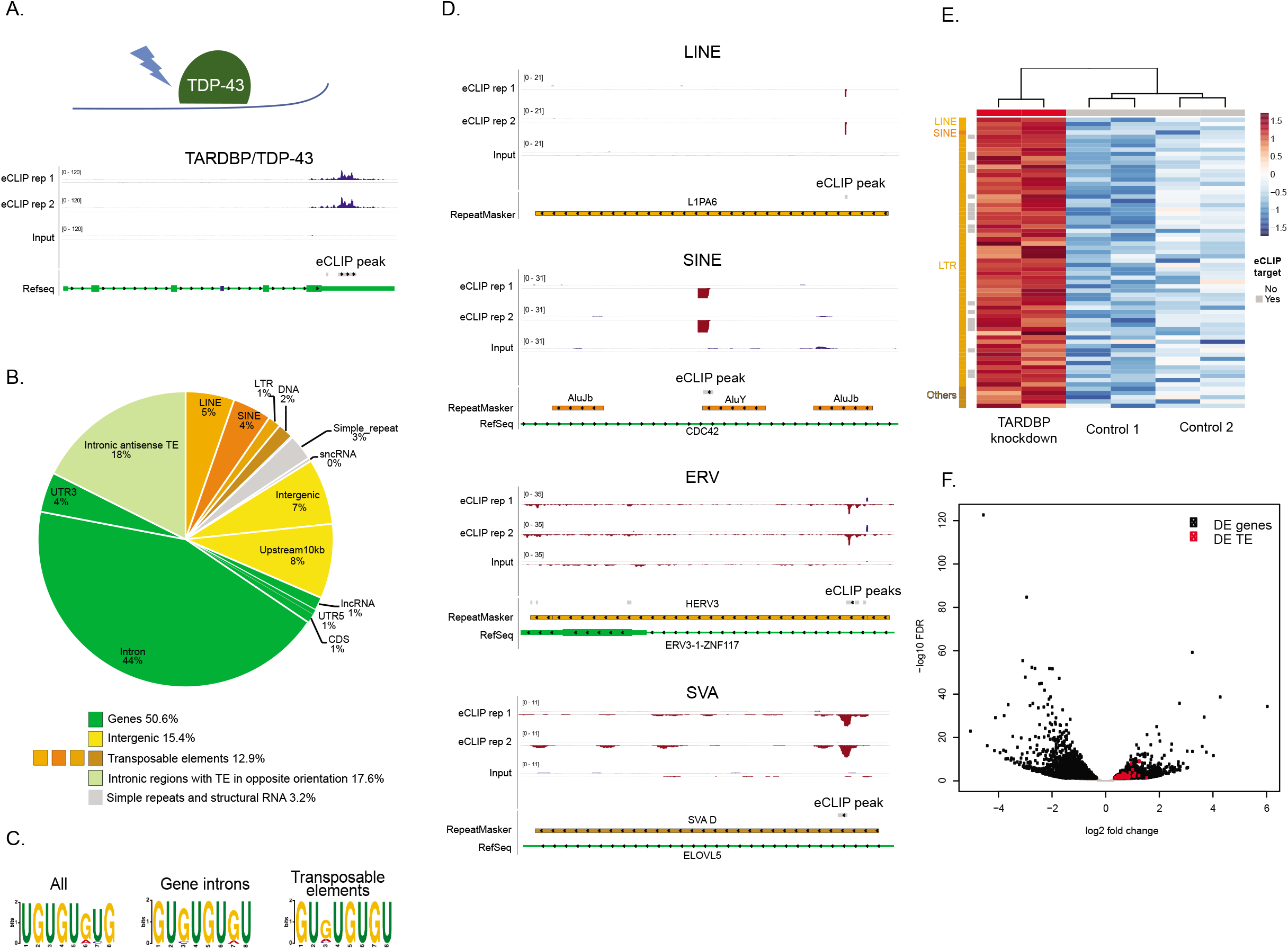
TDP-43 binds and regulates retrotransposon transcripts. (**A**) A schematic of the eCLIP-seq based identification of directly bound RNA targets for TDP-43, with a representative pileup of reads where TDP-43 is known to target its own 3’ UTR. (**B**) A pie chart of annotation categories for TDP-43 targets shows that retrotransposons are a small but important fraction of direct targets (12.9%). Gene intronic sequences represent the largest fraction of all peaks at 44%, while an additional 17.6% represent regions where TEs provide regulatory sequences to the host gene intron. (**C**) The known TDP-43 binding motif (UGUGU) is present in both gene and retrotransposon targets. (**D**) Representative browser pileup plots for one retrotransposons from each major repeat class: an L1PA6 LINE element, an AluY SINE element, a HERV3 LTR element, and an SVA_D from the human specific SVA class. (**E**) Knockdown of TDP-43 levels results in the upregulation of retrotransposon transcripts, most of which also show evidence for direct binding in the eCLIP-seq experiments (marked by grey bars at the head of each row). (**F**) A volcano plot of fold change versus FDR q-values shows that up-regulated TEs (red dots) form a substantial fraction of all TDP-43 dependent genes, and all are up-regulated.

To determine the subset of CLIP targets that are regulated at the transcript abundance level, we next knocked down the levels of TDP-43 protein using a short hairpin RNA strategy (TDP-43kd), and sequenced the transcriptomes of both control and TDP-43kd libraries. All significantly altered retrotransposon transcripts were up-regulated in TDP-43kd cells, indicating that TDP-43 normally contributes to the silencing of retrotransposon transcripts (Fig 5E, Supp. Table S11). This is in contrast to altered gene transcripts, where a substantial fraction was either up- (1165 genes, 34%) or down-regulated (2246 genes, 66%) in TDP-43kd cells (See Fig 5F and Supp. Table S11). Moreover, a majority of the directly bound gene targets were down-regulated in the absence of TDP-43 (52%). This is consistent with known roles for TDP-43 in contributing to alternative splicing or translation control for a large fraction of gene targets that are not regulated at the mRNA abundance level (Polymenidou et al., 2011; Tollervey et al., 2011). However, the fact that nearly all expressed retrotransposon transcripts were up-regulated in the absence of TDP-43 suggests that TDP-43 plays a silencing role for transposons. This is consistent with previous studies of the role of TDP-43 in the fly brain (Krug et al., 2017), but this is the first study to conclusively link directly bound TDP-43 transposon targets with the regulation of these targets at the expression level.

### C9orf72 repeat expansions are not associated with ALS subtype

A hexanucleotide repeat expansion in *C9orf72* is the most common pathogenic alteration associated with the development of ALS, both familial and sporadic (Majounie et al., 2012). Moreover, previous studies have noticed a slight trend toward higher TE expression in patients carrying C9orf72 mutations (Pereira et al., 2018; Prudencio et al., 2017; Y.-J. Zhang et al., 2019). In order to determine if patients carrying a *C9orf72* repeat expansion are overrepresented in one or several of the identified ALS subtypes, genotyping of the hexanucleotide repeat size was performed on the NYGC patient samples. *C9orf72* repeat expansion carriers were neither significantly over-nor underrepresented in the ALS-TE samples (p<0.55). However, we observed a nonsignificant trend towards enrichment of *C9orf72* repeat expansion in the ALS-Glia samples (p<0.08), and a slight depletion in the ALS-Ox samples (p<0.05). We note that none of the patient samples from the UCSD validation cohort carried repeat expansions in C9orf72, though each of the ALS subtypes was represented in this group (Supp Fig S2) Together this suggests samples from patients with *C9orf72* repeat expansions are not driving the transcriptional profiles of these subgroups, and show no particular enrichment in the ALS-TE group.

## Discussion

*De novo* discovery of transcriptome subtypes from a large patient sequencing study demonstrated that ALS patients fall into three distinct subtypes. The pathways that define two of these subtypes (oxidative & proteotoxic stress and neuroinflammation) have a well-established association with ALS disease, while a third subtype showed highly elevated retrotransposon expression. Several previous studies have described roles for neuronal stress and neuroinflammation as contributing factors for ALS. While genetic mutations are rare in ALS disease, the particular gene pathways and molecular processes altered by these mutations have allowed for insights into the mechanistic basis for ALS development and progression. This has led to a general grouping of the ALS-associated genes into those that largely mediate protein homeostasis, RNA metabolism, and neuroinflammation (Taylor et al., 2016), though frequently each gene can be linked to more than one of these processes. An example is SOD1, which has been ascribed roles in oxidative stress (Rosen et al., 1993), proteotoxic stress (Bruijn et al., 1998), and microglial activation (Chiu et al., 2013). While multiple cellular stressors might be contributing to the disease, we found that each patient sample could be grouped into distinct subtypes, which largely reflected pathways already described for ALS, but which fell into surprisingly distinct groups. For example, 60% of the patient samples exhibited gene expression markers suggesting oxidative stress and proteotoxic stress were the main contributors to cellular dysfunction (the ALS-Ox group), but did not show elevated markers of glial cell types or inferred glial cell enrichment in the tissue downstream of these stress responses. Only 20% of the patients showed extensive glial involvement (the ALS-Glial group), with an unknown mechanism initiating the activation of glial cells for these patients in particular.

The role of transposable element expression is a relatively new topic in the study of neurodegeneration. Moreover, the connection between TDP-43 and transposable elements has only recently been explored. While previous studies have linked TDP-43 to transposon binding in human cells and to transposon regulation in animal models (Krug et al., 2017; Wenxue Li et al., 2015), this is the first to link TDP-43 bound transposon transcripts to those that are de-silenced in human cells and to demonstrate that these same targets are de-silenced in a subset of human ALS patients. Additional mechanisms for transposon de-silencing in ALS patients with C9orf72 repeat expansions have been suggested (Pereira et al., 2018; Prudencio et al., 2017; Y.-J. Zhang et al., 2019), though we note that C9orf72 status was not associated with the ALS-TE group in this study. Together, this suggests a model where retrotransposon silencing is a normal function of TDP-43 in somatic cells, and that this role is disrupted in ALS patient tissues, potentially contributing to cellular toxicity. Previous studies have demonstrated that expression of envelope proteins from the endogenous retrovirus class of retrotransposons can be toxic to cells of the central nervous system (Antony et al., 2004; Kremer et al., 2013; Wenxue Li et al., 2015). Additional studies have shown that expressed retrotransposon RNAs can also be toxic through aberrant recognition by innate immune components in Aicardi-Goutieres Syndrome (Crow and Manel, 2015). Other studies linked transposon activation in Alzheimer’s disease with tau aggregation, and suggest these alterations accompany neuroinflammation and genomic instability (Guo et al., 2018; Sun et al., 2018). DNA damage and structural variants induced by transposition itself are a formal possibility, given the elevated levels of the fully competent LINE-1 Hs retrotransposon, and evidence for active L1Hs transposition in human neurons (Evrony et al., 2016; Upton et al., 2015). However, most of the retrotransposons expressed in this study derived from fixed elements that have lost the capacity to transpose. These and other mechanisms for retrotransposon contributions to cellular toxicity remain to be explored in ALS patients, but present a possible mechanism for cellular damage in the subset of patients with extensive TDP-43 pathology and retrotransposon re-activation.

Finally, the three subgroups defined in this study were identified from a largely sporadic set of patients with no known family history of the disease. Thus, ALS subtype appears to be largely independent from genotype. Transcriptional differences that separate these subtypes may reflect multiple contributing factors including causal mechanisms, disease progression stage, cellular composition of the tissues due to loss of particular cell types, and differences in environmental interactions that contribute to onset. The fact that transcriptional profiles of post mortem cortex tissues segregate into three distinct groupings may provide an entry point to investigate these underlying mechanisms. If peripheral tissues amenable to sampling or biopsy (e.g. peripheral blood monocytes or muscle) have distinct molecular signatures corresponding to these CNS groups, it could facilitate cohort selection for clinical trials of therapies targeting specific pathogenic mechanisms. Indeed, the existence of distinct clusters of sporadic ALS patients – whether based on intrinsic pathogenic differences or disease stage, might explain why many treatments identified in laboratory models targeting a specific pathway have failed to translate to successful clinical trials. Additional correlations may arise as the ALS Consortium cohort size grows in terms of number of patients profiled, tissues examined, and data types collected.

## Materials and Methods

### Acquisition of patient materials

UCSD samples were obtained from patients who had been followed during the clinical course of their illness and met El Escorial criteria for definite ALS. These individuals had bulbar or arm onset of disease and caudally progressing disease. Control nervous systems were from patients from the hospital’s critical care unit when life support was withdrawn. All samples were genotyped for C9orf72 repeat expansions using standard repeat-primed PCR, as previously described (Renton et al., 2011). Briefly, primers outside the C9orf72 intronic repeats were amplified in a nested PCR strategy, and the resulting traces were analyzed on a Bioanalyzer to determine repeat length.

The NYGC ALS Consortium samples, the majority provided by the Target ALS Post-Mortem Tissue Core, were acquired through various IRB protocols from member sites and transferred to NYGC in accordance with all applicable foreign, domestic, federal, state, and local laws and regulations for processing, sequencing, and analyses. Many of these samples were genotyped for C9orf72 repeat expansions using a combination of cross-repeat fragment size analysis, standard repeat-primed PCR, and the Asuragen AmplideX PCR/CE C9ORF72 Kit. All available de-identified clinical and pathological records were collected and used together with C9orf72 genotypes to summarize patient demographics and disease features.

### Generation of RNA-seq libraries from UCSD ALS samples

RNA was extracted from each of the flash-frozen patient samples using the Ambion PureLink RNA Mini kit. RNA was assessed using the Bioanalyzer to ensure RNA Integrity (RIN) values of > 5.5. RNA-seq libraries were prepared from 500 ng of total RNA using the Illumina TruSeq Stranded Total RNA kit. Samples were barcoded to multiplex 8 samples per batch, randomly mixing ALS and Control samples in each library preparation and sequencing batch. The libraries were sequenced on an Illumina NextSeq using a single end 76 nucleotide setting, and pooled to ensure ~40 million reads per library.

### Generation of RNA-seq libraries from NYGC ALS samples

RNA was extracted from flash-frozen patient samples homogenized in Trizol-Chloroform and purified using the Qiagen RNeasy Mini kit. RNA was assessed using the Bioanalyzer. RNA-seq libraries were prepared from 500 ng of total RNA using the KAPA Stranded RNA-seq kit with RiboErase for rRNA depletion and Illumina compatible indexes (NEXTflex RNA-seq Barcodes, BioScientific). Pooled libraries (average insert size: 375bp) were sequenced on an Illumina HiSeq 2500 using a paired end 125 nucleotide setting, to yield ~40-50 million reads per library.

### shRNA knockdown and RNA-seq in SH-SY5Y cells

SH-SY5Y cells (CRL-2266, ATCC) were grown in DMEM/F-12 Dulbecco’s Modified Eagle Medium/Nutrient Mixture F-12 (ThermoFisher, 11320033) supplemented with 10% FBS and 1% penicillin–streptomycin, and cultured at 37□°C with 5% CO_2_.

The MISSION pLKO.1-puro human TDP-43 (TRCN0000016038) and control shRNAs (Sigma Aldrich SHC007 and SHC016) were used to produce lentivirus. The pLKO.1 plasmid DNA, together with psPAX2 packaging (Addgene, 12260) and pMD2.G envelope (Addgene, 12259) plasmid DNA were combined at a ratio of 4:3:1, respectively and transfected with PEI reagent (Polysciences) into HEK293FT cells. Virus containing medium was collected 48 and 72 hours post-transfection and stored at −80°C.

One and a half to two million SH-SY5Y cells per well of a 6-well plate were “spinfected” with 3 mL of virus containing media and 8μg/mL polybrene for 1 hour at 800g. Cells stably expressing shRNAs were selected with 2 μg/mL of puromycin 48 hours postinfection for 3 days. Uninfected cells at similar density were used as a control for puromycin selection. Cells were harvested 5 days after infection for downstream analyses.

Total RNA was extracted using Trizol (ThermoFisher) according to the manufacturer’s instructions. One microgram of total RNA was subjected to rRNA removal using the Ribo-Zero Gold rRNA Removal kit (Illumina). Strand specific RNA-seq libraries were constructed using NEBNext Ultra II Directional Library Prep Kit (NEB). The libraries were sequenced on an Illumina NextSeq using a single end 76 nucleotide setting, and pooled to ensure ~40 million reads per library.

### TARDBP eCLIP in SH-SY5Y cells

Twenty million SH-SY5Y cells were UV-crosslinked (254 nm, 400 mJ/cm2) in PBS and cell pellets were stored at −80°C. Each sample was prepared as an independent biological replicate and included non-crosslinked cells as a control. Cell pellets were lysed in 1 mL lysis buffer, followed by RNAse I digestion and immunoprecipitation with 10 μg of anti-TDP-43 antibody (10782-2-AP, Proteintech). Precipitated RNA was reverse-transcribed using SuperScript IV (Thermo Fisher Scientific) and eCLIP libraries were prepared as previously described (Van Nostrand et al., 2016). The libraries were sequenced on an Illumina NextSeq500 using a single end 76 nucleotide setting.

### Histology and Immunohistochemistry

Frozen motor cortex tissues were fixed in 10% neutral buffered formalin and 5 μm paraffin sections were cut for Hematoxylin and Eosin (H&E) and immunohistochemistry (IHC) staining. IHC staining was performed on the Ventana Discovery Ultra platform with OmniMap HRP and ChromoMap DAB detection system according to manufacturer’s protocols (Roche), using primary antibodies specific for TDP-43 (10782-2-AP, Proteintech, Rosemont, IL, USA, 1:10,000) and phosphoTDP-43 (pS409/410) (CAC-TIP-PTD-M01, Cosmo Bio USA, Co., Carlsbad, cA, 1:200). Stained slides were scanned by Leica Aperio ScanScope system.

### Analysis of RNA-seq libraries

Reads from samples with RIN >= 5.5 were aligned to the hg19 human genome using STAR v2.5.2b (Dobin et al., 2013), allowing for a 4% mismatch rate and up to 100 alignments per read to ensure capture of young transposon sequences. Abundance of gene and transposon sequences was calculated with TEtranscripts v2.0.3 (Jin et al., 2015). For differential expression analysis, we employed DESeq2 (Love et al., 2014), using the DESeq normalization strategy, and using an FDR-corrected P-value threshold of P<0.05. For heatmap visualization, the reads were normalized using a variance stabilizing transformation in DESeq2.

### Transcriptome De Novo Cluster Identification

The number of clusters in the ALS datasets was determined using the SAKE software suite (Ho et al., 2018). Briefly, a variance stabilizing transformation was performed on the raw counts data using DESeq2. Gender associated genes were then removed from this list before rank ordering by median absolute deviation using the SAKE software suite, selecting the top 5000 genes with the highest MAD. SAKE was implemented using 200 iterations, the “nsNMF” algorithm for Non-negative Matrix Factorization, and a “k” setting of 3 clusters for the cortex samples, determined by the k setting with the highest cophenetic correlation coefficient.

### Classification of UCSD samples using NYGC ALS subtypes

A variance stabilizing transformation was performed on the raw counts of NYGC and UCSD data using DESeq2. Principal component analysis was then performed to identify principal components that delineate the identified ALS subtypes, and a 95% confidence ellipse was calculated from the NYGC samples for each subtype. The position of the UCSD patient samples along the principal components of interest was determined as the midpoint of the two replicates, and a distance to the center of the 95% confidence ellipse for each subtype was calculated. The UCSD samples were classified based on the nearest ellipse center of the subtype.

### Analysis of eCLIP libraries

Reads were trimmed to remove adapters, and aligned to the hg19 human genome using STAR v2.5.2b (Dobin et al., 2013), allowing for a 4% mismatch rate and up to 100 alignments per read to ensure capture of young transposon sequences. Weighting of multi-mapper alignments (expectation-maximization modeling) and identification of regions of enrichment relative to input (using negative binomial modeling) were performed with CLAM v1.1.3 (Z. Zhang and Xing, 2017). Integrated genome viewer (Robinson et al., 2011) was used for visualization of enriched regions and read depth.

### Gene Set Enrichment Analysis

Pre-generated gene sets used for GSEA (Subramanian et al., 2005) were obtained from the MSigDB Hallmark collection (Liberzon et al., 2015), and KEGG (Kanehisa and Goto, 2000). Custom gene sets were generated from the SOD1(Chiu et al., 2013) and TREM2 (Keren-Shaul et al., 2017) models of disease-associated microglia, as well as TARDBP targets from this study. Custom transposable element sets, including retrotransposons considered active in the human genome (Mills et al., 2007) were generated from RepBase (Bao et al., 2015).

### Statistical Analysis

All RNA-seq analyses employed negative binomial modeling and B-H corrected FDR P-values via the DESeq software (Love et al., 2014). All CLIP-seq analyses employed negative binomial modeling and B-H corrected FDR P-values via the CLAM software (Z. Zhang and Xing, 2017). Differences between ALS Subgroups and correlations with gene expression or cell type composition used Wilcoxon Mann Whitney U-tests. Enrichments for genotype in each subtype used Fisher’s Exact test.

### Data Availability

All data from this study outside of the ALS Consortium datasets have been deposited at the following GEO accession: GSE122650. RNAseq data generated through the NYGC ALS Consortium in this study can be accessed at GEO accession: GSE124439.

## Acknowledgments

We wish to thank Y Jin for helpful discussions, the Target ALS Human Post-Mortem Tissue Core for providing post-mortem brain samples, the CSHL Sequencing Facility (supported by an NIH Cancer Center Support Grant 5P30CA045508) for additional sequencing support, the CSHL Histology Core Facility (partially supported by NIH Support Grant 5P30CA045508) for performing the histology and IHC staining, and the Harms laboratory for providing C9ORF72 genotypes for the Target ALS samples. Schematic images were adapted with permission from Servier Medical Art (https://smart.servier.com). This work was supported by the Ride For Life Foundation, the Rita Allen Foundation of which MGH is a scholar, a Ben Barres Early Career Acceleration Award from the Chan Zuckerberg Initiative to MGH, the O’Neil Charitable Trust, the NIH/NINDS through grants: 5R21NS088449 and R01NS091748, and the NIH/NIA through grant: R01AG057338-01. All NYGC ALS Consortium activities are supported by the ALS Association (grant number 15-LGCA-234) and the Tow Foundation.

## Author contributions

OHT, MGH and JD designed the study. NVR designed and performed the experiments identifying TDP-43 targets in SH-SY5Y cells. RS designed and performed the experiments on the UCSD ALS patient samples. JR provided the UCSD ALS patient samples and associated clinical and diagnostic data. In the NYGC ALS Consortium, members contributed ALS patient samples and clinical information. DK curated deidentified clinical data and C9orf72 genotype information; IH and NP coordinated study materials and processed samples for sequencing; SF oversees Consortium resources and data distribution. DF and HP designed the methodology, reviewed sample preparation and data quality, and coordinated the research activity of NYGC ALS Consortium postmortem core RNAseq experiments. LWO coordinated ALS postmortem sample and data collection through the Target ALS Multicenter Post-Mortem Tissue Core. OHT and MGH analyzed the data. All authors contributed to the interpretation, writing and editing of the manuscript.

## Materials & Correspondence

All correspondence and requests for materials should be addressed to: M. Gale Hammell.

## Supplementary Materials

**Supplementary Fig. 1. Determination of optimal cluster numbers by SAKE for de novo clustering of the NYGC transcriptomes. (A)** Consensus clustering heatmaps to measure sample correlation distances for a range of values “k” of clusters present from 2-5. **(B)** Statistical measures of cluster consistency for each of the estimated cluster assignments in (A). **(C)** A heatmap depicting the relative confidence of cluster assignments for each sample in the NYGC transcriptome for k=3 clusters. **(D)** A heatmap depicting the relative confidence in each gene marker in determining cluster assignments for k=3 clusters. **(E)** A consensus clustering heatmap depicting within cluster distances for k=3 clusters for a larger set of 1000 iterations. **(F)** Scatterplot of the first two principal components (PC1 and PC2) showing distribution of the SAKE identified clusters for ALS-TE (red), ALS-Ox (Blue), and ALS-Glia (yellow).

**Supplementary Fig. 2. Description of the motor cortex samples obtained from the UCSD tissue bank. (A)** Schematic description of tissues obtained for 13 sporadic ALS patients and 6 non-neurological controls. **(B)** PCA plot showing the position of UCSD samples (marked by JRxx patient ID numbers) relative to the subgroups within the NYGC ALS Consortium cohort. Samples assigned to the ALS-TE group (red), ALS-Ox group (blue), and ALS-Glia group (gold) fall largely within the 95% confidence interval ellipses defined by the ALS Consortium samples. **(C)** Heatmap showing the expression patterns for the ALS subtype markers for the NYGC and UCSD samples.

**Supplementary Fig. 3. NeuroExpresso based marker gene profile (MGP) signatures of additional cortical cell types among the ALS subtype samples**. Estimation of astrocyte, oligodendrocyte, microglia (resting and activated), GABA-ergic neuron, pyramidal neuron and endothelial cell composition distributions within the ALS subtypes and controls. Benjamini-Hochberg adjusted P values less than 0.05 (when compared to control) are indicated.

**Supplementary Fig. 4. TDP-43 and pTDP-43 histology analysis**. Example sections for tissue slices stained for TDP-43 and pTDP-43 are shown for UCSD patients assigned to the ALS-TE group (JR34, JR47), the ALS-Ox group (JR9), the ALS-Glia group (JR27), and two non-neurological controls (JR65, JR67). Asterisks mark example cells with large pTDP-43 positive inclusion bodies in the ALS-TE samples, which were absent from the tissues for each of the other patients.

**Supplementary Table 1. Clinical information for the NYGC ALS Consortium samples**

**Supplementary Table 2. Full list of SAKE-identified features marking each of the ALS subtypes in the NYGC ALS Consortium patient cohort**.

**Supplementary Table 3. GSEA gene sets with FDR < 0.05 in ALS-Ox subtype**

**Supplementary Table 4. GSEA gene sets with FDR < 0.05 in ALS-Glia subtype**

**Supplementary Table 5. GSEA gene sets with FDR < 0.05 in ALS-TE subtype**

**Supplementary Table 6. NYGC ALS consortium patient subtype assignment**

**Supplementary Table 7. Clinical information for the UCSD sALS samples**

**Supplementary Table 8. List of genic targets bound by TDP-43 eCLIP**.

**Supplementary Table 9. List of transposable elements and satellites bound by TDP-43 eCLIP**.

**Supplementary Table 10. Enriched motifs identified under the TDP-43 eCLIP peaks**

**Supplementary Table 11. Differentially expressed genes and transposable elements upon TDP-43 knockdown**

**Supplementary Table 12. Principal investigators of the NYGC ALS Consortium**

